# A fast, scalable, MinHash-based k-mer tool to assess Sequence Read Archive next generation sequence submissions

**DOI:** 10.1101/2021.02.16.431451

**Authors:** Kenneth S. Katz, Oleg Shutov, Richard Lapoint, Michael Kimelman, J. Rodney Brister, Christopher O’Sullivan

## Abstract

Sequence Read Archive submissions to the National Center for Biotechnology Information often lack useful metadata, which limits the utility of these submissions. We describe a scalable k-mer based tool for fast assessment of taxonomic diversity intrinsic to submissions, independent of metadata. We show that our MinHash-based k-mer tool is accurate and scalable, offering reliable criteria for efficient selection of data for further analysis by the scientific community, at once validating submissions while also augmenting sample metadata with reliable, searchable, taxonomic terms.

## Background

Established in 2007, the National Center for Biotechnology Information (NCBI) Sequence Read Archive (SRA) accepts raw sequencing data directly from high-throughput sequencing platforms (1). Next generation sequencing (NGS) sets are inherently large, and improved technologies are exquisitely sensitive to contamination. Submissions must be processed, before either interpretation or quality assessment is possible, to provide submitter feedback and submission verification. The growth of data submission is exponential (doubling approximately every 12 months (2)), rendering use of computationally expensive methods, such as *de novo* assembly followed by alignment, impractical due to costs and limits of scale, particularly given the time constraint of submission processing.

We considered that questions about the quality of a given NGS run could reasonably be inferred from the taxonomic distribution of reads within that set, whether based on a single organism or of metagenomic design. This is often enough information to answer basic experimental or clinical questions, as well as inform decisions about the merit of subsequent resource-intensive assessment methods. Read sets with organismal tags can be used to select data for further analysis. Moreover, binning reads into taxonomic buckets can identify contaminating reads and reads outside of the stated experimental scope. Such identified reads can be filtered from a sample before downstream processing. This proposed taxonomic analysis is independent of metadata and intrinsic to the run, capable of both validating submissions and augmenting sample metadata with reliable, searchable, taxonomic terms.

Following these principles, we developed a k-mer-based Sequence Taxonomic Analysis Tool (STAT). Based on MinHash (3), and inspired by Mash (4), STAT employs a reference k-mer database built from available sequenced organisms to allow mapping of query reads to the NCBI taxonomic hierarchy (5). We use the MinHash principle to compress the representative taxonomic sequences by orders of magnitude into a k-mer database, followed by a process that yields a set of diagnostic k-mer hashes for each organism. This allows for significant coverage of taxa with a minimal set of diagnostic k-mers. Our results show STAT is a reliable method for examining submitted NGS data in a timely, and scalable, manner.

## Results

STAT was developed for quality assessment of SRA submissions to be shared with the submitter, requiring that analyses ideally take no more time than that of existing submission processing, while minimizing resource usage. Our design starts from the MinHash principle that a random selection of the lowest valued constituent blocks in a pool after hashing represents a signature of the parent object. In building k-mer databases from the set of sequences assigned a specific NCBI taxonomy id (TaxId), we read the 32 base pair (bp) k-mers as 64-bit hashes, selecting the minimum value representative for a window, then iteratively merging k-mers from taxonomic leaves to roots (see Methods, Figures 1, 2).

**Figure 1.**
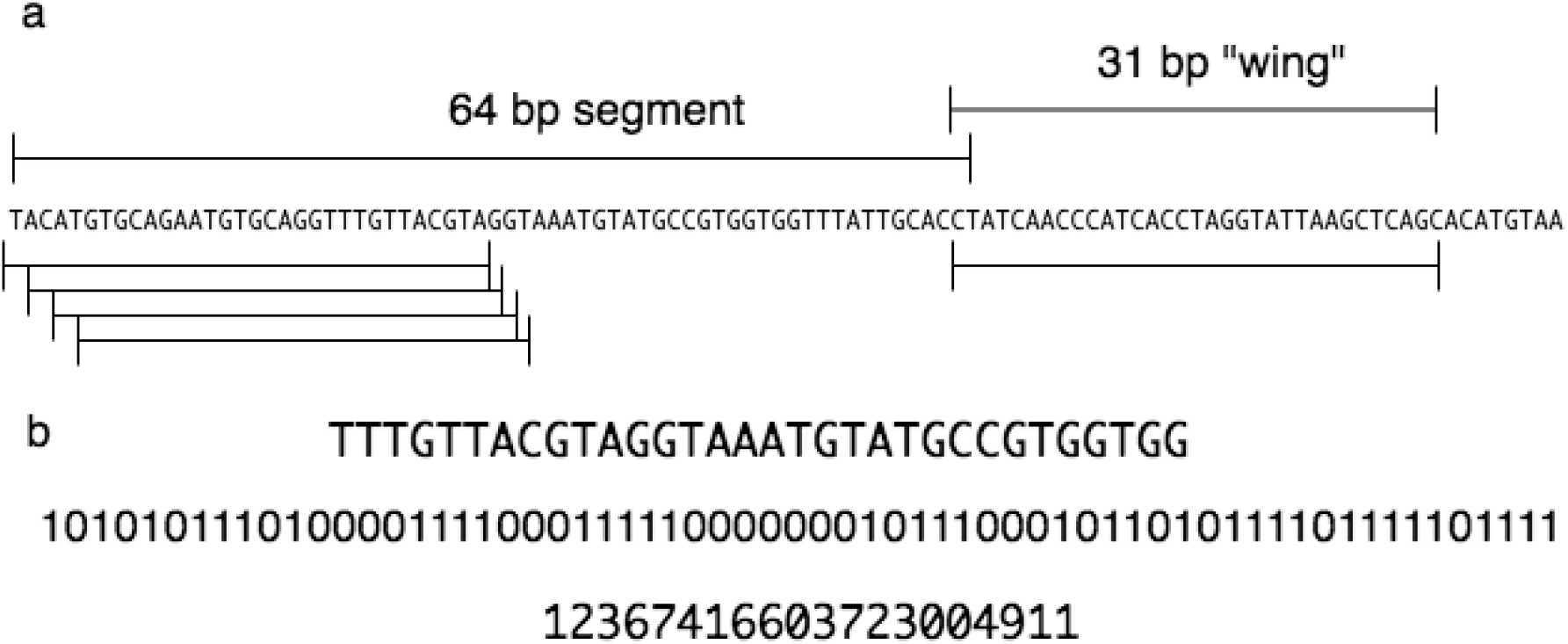
*Finding a minimum representative 32 base pair k-mer*. **a** From a selected 64 base pair segment, the series of 64 possible 32 base-pairs k-mers is defined by sequentially shifting the 32-base window by one base. The first four, and last of the possible k-mers is shown schematically. **b** An example selected k-mer sequence is shown followed by: first the two-bit encoding of that same k-mer sequence; finally the 64-bit decimal value of the encoded k-mer. The lowest 64-bit decimal value is selected as the representative k-mer for this 64 base segment.

**Figure 2.**
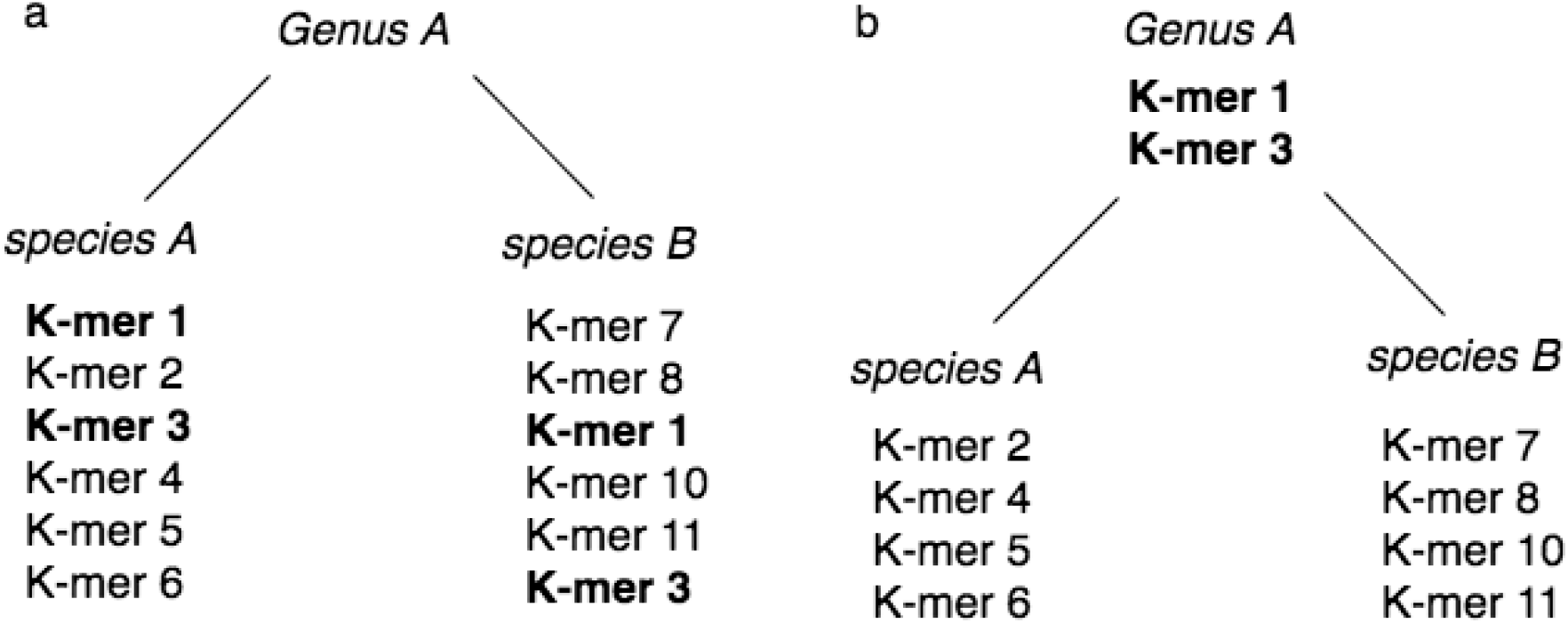
*K-mer taxonomic merging*. See Methods for details. **a** Before merging two sibling species are depicted containing both unique and shared k-mers (indicated in bold).. **b** After merging the two shared k-mers “merge” up to the *genus* level, while those unique to each remain diagnostic for the *species*.

Initial analysis using only densely populated k-mer databases performed well. However, despite being on average over an order of magnitude smaller than the input sequence database size (see below), we determined that loading the entire densely merged “tree_filter.dbs” into memory for analysis unnecessarily incurred long I/O read time and large memory costs since most runs required only a fraction of the complete database. Moreover, STAT jobs, like many computational pipelines, are submitted to either a local computer farm cluster scheduler (“grid engine”), or by dispatching cloud-based virtual machines. In both cases job scheduling typically requires explicit needed resource declarations such as CPU and memory. An initial screen capable of evaluating diversity of the sample and necessary resource requirements for detailed analysis minimizes cost and maximizes computational efficiency. For these reasons we pursued a selective two-step analysis, using a sparse filtering database in the first step to identify the presence of any (a) eukaryote if there are more than 100 biological reads of a species, (b) bacteria, or archaea with more than 10 biological reads, and (c) virus if there are 1-2 biological reads. This first pass is neither qualitative, nor exhaustive, but allows us to quickly identify taxa for focus in the second pass (Figure 3).

**Figure 3.**
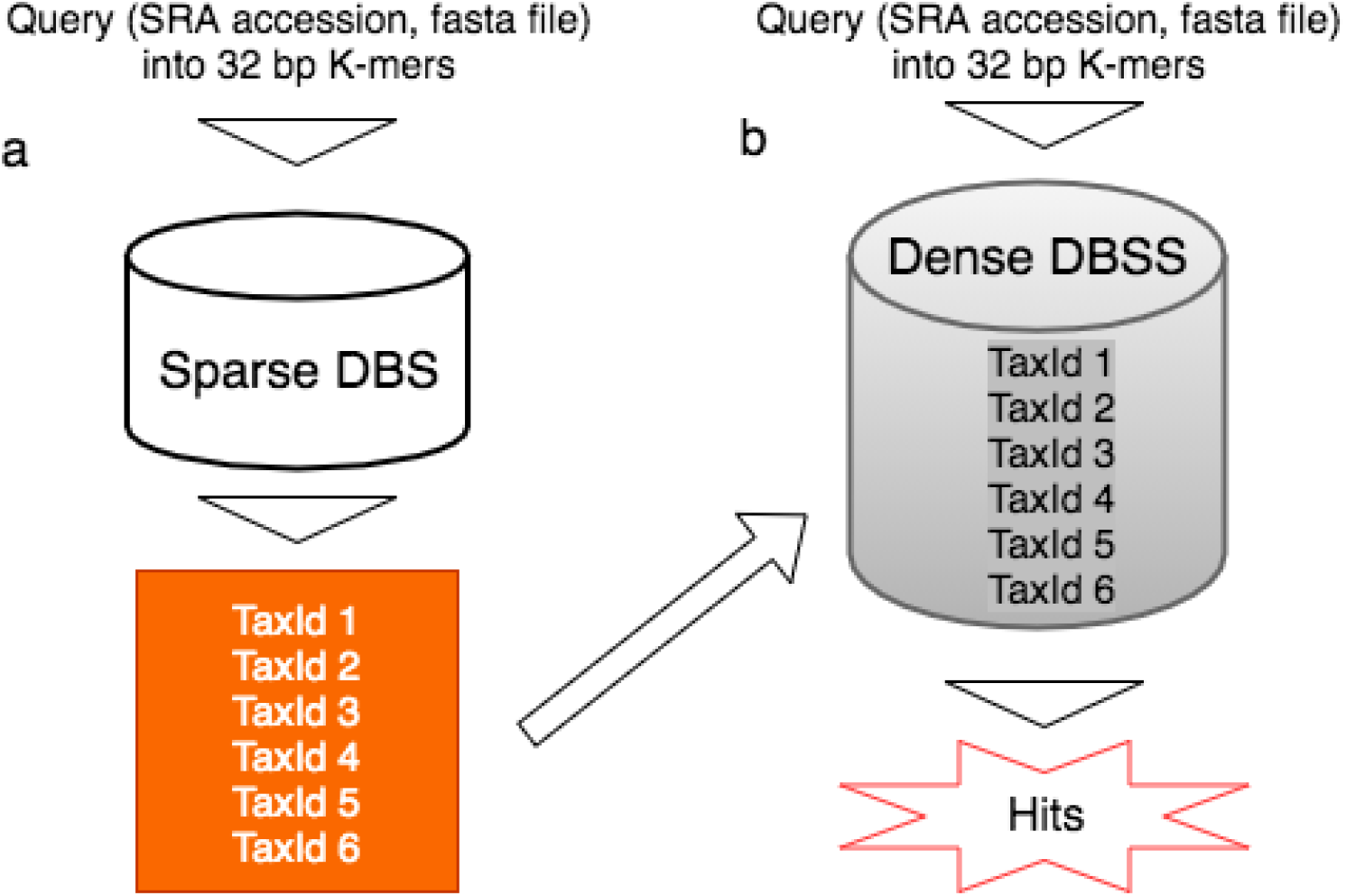
*STAT two phase query*. **a** In the first qualitative phase the input query (an SRA accession, or fasta file) is sequentially rendered into 32 bp k-mers, and matches to the decimal hash values found in the sparse database identifying taxa for deeper analysis. **b** The TaxIds identified are used to compose the dense database using the same query in the second quantitative pass.

To facilitate this two-step process, and further minimize resource requirements, we decreased k-mer database size by 33% by storing only the 8-byte k-mers in a database file, separately storing pairs of TaxId, total TaxId k-mer count for each TaxId respectively in an auxiliary “annotation” file. The k-mer database / k-mer count annotation file pair is designated “dbss”, the database sorted by TaxId, with each TaxId set sorted by k-mer. TaxIds identified in the first step against the sparse k-mer database are used in the second step to load into memory only those TaxId k-mer hashes using the counts provided by the annotation file as offsets. MinHash sampling combined with dynamic loading of only necessary dense TaxId database k-mer hashes yields significant benefits for cpu and memory requirements. Further, the selection of TaxIds to load may be augmented by heuristics, such as purposely withholding TaxIds from contamination detected in the prior filtering step.

STAT reports the distribution of biological reads mapping to specific taxonomic nodes as a percentage of total biological reads mapped within the analyzed run. Since results are proportional to the size of sequenced genomes, a mixed sample containing several organisms at equal copy number is expected to find more reads originating from the larger genomes. This means that percentages reported likely reflect sample genome size(s), and must be considered by the user against the genomic complexity of the sequenced sample.

Like all sequence-based classification schemes, STAT reflects and depends upon accurately encapsulating evolutionary paths. The significant achievements of adapting both the NCBI RefSeq data model (6), and internationally accepted taxonomy to incorporate metagenomic viral sequence (7,8) fundamentally benefit STAT and other similar classification tools.

An important consequence of merging in k-mer database construction is to avert complications caused by biological complexities. For example, most k-mers derived from endogenous retroviruses found in the human input reference genome will likely merge to the root as those k-mers would also be found in the Viruses Super Kingdom.

Further, when analyzing results each level – read, run – requires integration of less than ideal signals. It is common to find multiple TaxIds identified in a single biological read, ideally coherent for a given lineage. Were those *Mus musculus*, Murinae, and Mammal, there is confidence in declaring the read *Mus musculus*. Should a read map to multiple, related taxonomic nodes, it is reported as originating from the most proximal shared taxonomic node. For example, a read with hits to sibling species may be reported as their common genus, conservatively locating the most proximal common node before ambiguity (Figure 4). Likewise, such conservative heuristics are required when integrating the signals from all biological reads to report the run. If the run subject is a single organism, it is expected that STAT would identify taxonomic nodes across the lineage, and that the number of reads mapping to higher level nodes will be more than those mapping to terminal nodes.

**Figure 4.**
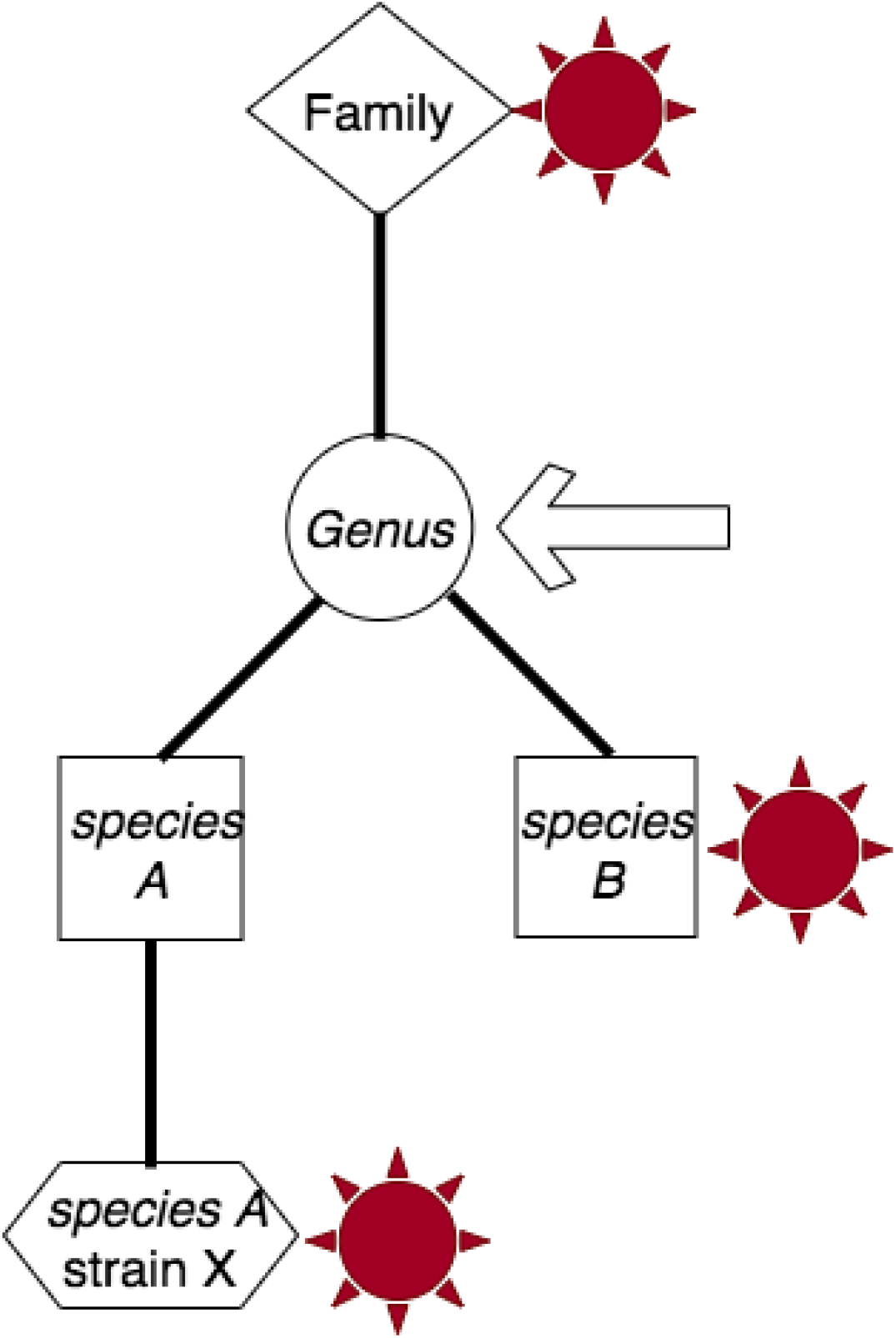
*Resolution of taxonomic assignment*. Depicted are three distinct k-mer hits within a single read to multiple branch taxa. The arrow indicates conservatively the most proximal unambiguous common node.

STAT was designed as a tool for assessing the quality of *any* SRA submission and has matured into a tool that also significantly enhances user comprehension. Many k-mer tools were created for the purpose of metagenomic taxonomic assignment (9) during STAT development. Taxonomic classifiers balance speed, accuracy, and memory requirements. While STAT was neither primarily developed for metagenomic analyses, nor as a tool for distribution, the same concerns apply. Using MinHash to sample and save at most 1 out of every 64 k-mers generated from input sequences yields k-mer databases 1-2 orders of magnitude smaller than the parental reference nucleotide database from which they were derived. For example, currently the BLAST^®^ refseq_genomes database used is 1.4 terabytes (tb) whereas the representative sparse and dense STAT k-mer databases are approximately 1.5 gigabytes (gb), and 75 gb, respectively.

The STAT k-mer databases contain 248,426 TaxIds before merging. Our complete merged 75 gb dense database (“tree_filter.dbss”) represents 130,817 TaxIds after merging (all data reflect the 20200518 build). Compare the Kraken default 70 gb database that only includes “RefSeq complete genomes, of which there are 2,256, while Kraken-GB contains 8,517 genomes” (10). Despite our sparse index database (“tree_index.dbs”) size of 1.5 gb, it nonetheless contains k-mers from 119,982 TaxIds.

We compare STAT accuracy to Kraken 2 using the strain exclusion test as described by Wood et al. (11). STAT shows the identical accuracy of Kraken 2 for both bacteria and virus (see Figure 5). As expected, STAT sensitivity is notably dampened as we chose to sample the widest taxonomic breadth. Our desire for conservative taxonomic assertions is further reflected by STAT never yielding a false positive in accuracy test results (data not shown).

**Figure 5.**
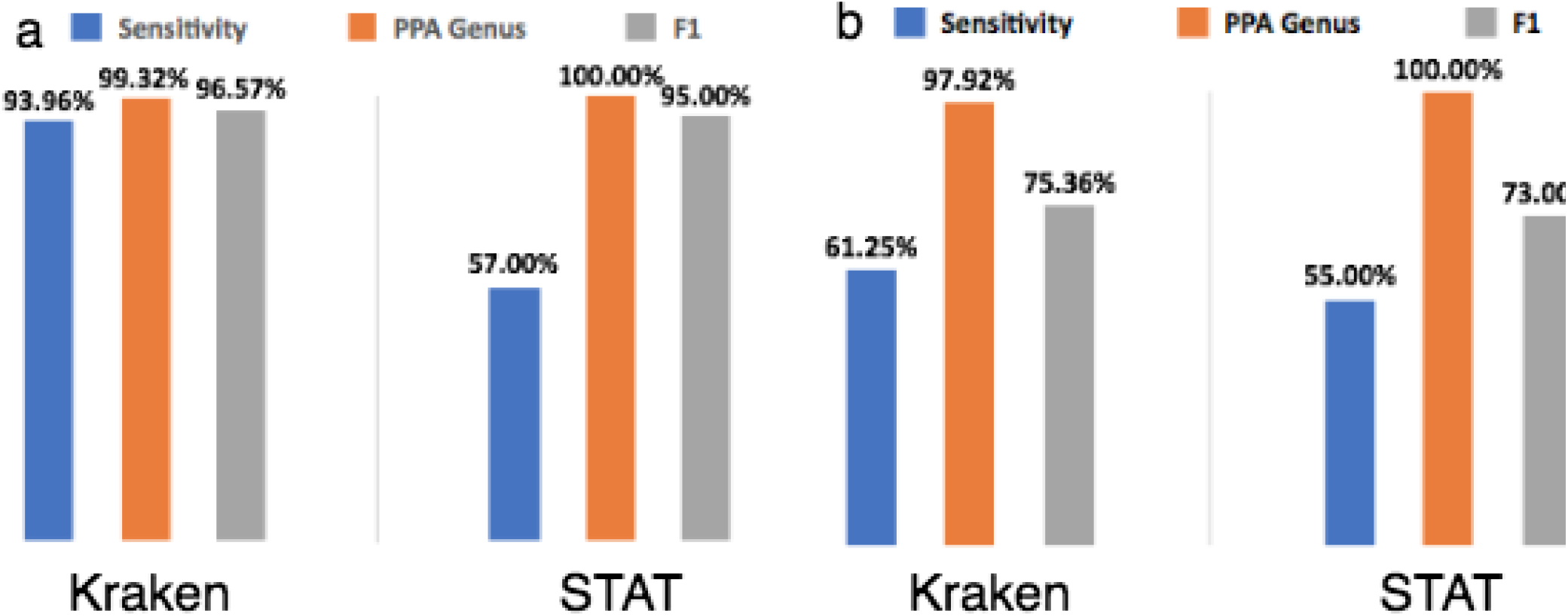
*STAT Accuracy and Sensitivity*. Comparison of STAT and *Kraken2* accuracy, sensitivity, and F1 measure using positive predictive value at the *genus* level for **a** Bacteria. **b** Virus.

We found it unnecessary to apply the same selection of k-mer hash minimums from query sequences to compose a similarity index (3,4), instead of exact k-mer (hash) matching. We show that accuracy is robust, while still reflecting our conservative bias in taxonomic assignment. Though similar in performance to Kraken 1 input speed (21.6 million reads/minute), and runtime (132.5 seconds) characteristics, STAT (maximum resident set size 830304 kilobytes) required only 8% and 1% of the memory needed by Kraken 1 and Kraken 2, respectively (11). Unsurprisingly, the accuracy test (see Methods) required additional time for extracting the requested TaxId k-mers on demand. Maximum resident set size during the accuracy test was approximately an order of magnitude greater than Kraken 2 (11, data not shown), despite loading a k-mer database 20 times the “strain_excluded” FASTA file size (3.9 gb) and over 100 times “strain_excluded.dbs” size (545 megabytes (mb)).

We provide two symmetrical examples of expected and unexpected contamination that illustrate STAT effectiveness.

### Contamination during a pandemic

Like many public health institutions worldwide Public Health England (PHE) programmatically surveils infectious pathogens using NGS, and submits targeted reference genomic analyses to SRA. The SARS-CoV-2 pandemic emerged in December 2019, and many countries outside China identified their first cases in early 2020 (12). The United Kingdom’s first cases were identified January 30, 2020 (13). Routine STAT analysis of submissions during early 2020 identified over 2,000 PHE surveillance bacterial NGS submissions likely contaminated with SARS-CoV-2 sequences. The earliest of these was dated February 11, 2020, less than two weeks from the first recognized U.K. cases. PHE was alerted to the likely carryover contamination, acting quickly to limit further events. Subsequent investigation confirmed SARS-CoV-2 contamination, ranging from a minimum of 1 positive spot^1^ containing 1 positive hit, reaching to 4,233 positive spots containing 18,270 hits (see Methods, and Additional file 1). This example underscores STAT utility in monitoring submissions for possible contamination, allowing curators to contact submitters to alert, and correct, the source of contamination.

### Identifying and removing potential personally identifying information

As lower cost significantly expanded human genome sequencing, awareness rose of potential personally identifying information residing in public repositories (14). Large efforts employing NGS to diagnose and monitor human health, or detect pathogenic outbreaks such as SARS-CoV-2, caused clinical sample submitters to worry about the inclusion of human sequence. As a counterpart to the previously discussed contamination example, we sought a STAT-based tool to find and remove unavoidable human sequence reads in clinical pathogen samples.

We began by building a k-mer database using human reference sequences withholding the iterative merging previously described. The majority (approximately 80%, see Methods) of k-mers derived represent conserved ancestral sequences, but our goal here is to aggressively identify human sequences. We then subtracted any k-mer also found in the merged kingdom databases Viruses, and Bacteria to protect against spurious false positive hits targeting clinical pathogens. After testing several window sizes, we found optimal performance using a segment of 32 bp (twice as dense as our standard taxonomy database).

Because unintended contamination is never uniform, we chose different ends of the expected spectrum of human content for testing (see Table 1). Two RNA_Seq runs were derived from bronchoalveolar lavage fluid taken from suspected SARS-CoV-2 patients. The wash of the lower respiratory tract from a patient suffering an active infection is expected to contain patient immune cells, sloughed patient epithelial cells, lung microbiota, and suspect clinical pathogens. Each run contains over five million spots, and though starting with approximately 85% Eukaryotic content, less than 10% of the spots remain after scrubbing for human sequence. The observation that a 3% selection of all possible human-derived 32 bp k-mers identifies over 92% of a random selection of likely human spots validates using MinHash and underscores its efficiency. These examples present a difficult test, and we identify 5-6% of the remaining spots as human.

**Table 1.** Summary of results including those found in Additional file 2 (S1-S4). We define “Human Spots” as those where all hits (up to top five) are identified as human with *eValue* < 1e -10. “Conserved Lineage Spots” are those containing a human top hit (lowest *eValue)* though not the exclusive organism of hits with *eValue* < 1e -10, and where all spot hits have either identical *eValue* or the greatest has *eValue* < 1e -14. “Total Length of Human Spot Alignments” is the sum of all the top alignments for all human spots maining.

|  | **Human RNA_Seq:**<br>**bronchoalveolar lavage fluid** |  | **SARS-CoV-2 Amplicon** |  |
| --- | --- | --- | --- | --- |
| **Accession** | SRR11092056 | SRR11092057 | SRR13402847 | SRR13444106 |
| **Total Spots** | 5239723 | 5184909 | 216859 | 471848 |
| **Total Spots**<br>**Remaining** | 438796 | 501436 | 216720 | 470934 |
| **Total Spots**<br>**Removed** | 4800927 | 4683473 | 139 | 914 |
| **Human Spots**<br>**Remaining** | 26265 | 25384 | 20 | 2 |
| **Conserved**<br>**Lineage Spots** | 27217 | 29507 | 70 | 13 |
| **Total Length**<br>**(kbp) of**<br>**Human Spot**<br>**Alignments** | 3684 | 3508 | < 3 | < 1 |

Unlike the previous examples, amplicon-directed sequencing of pathogens is expected to contain less unintended human content, as can be seen in Table 1. In both cases, 0.1% or less spots were removed, while among those remaining, 0.01% or fewer spots were identified as human. In no case was there any deleterious loss of the intended target signal (see Additional file 2, S5 Taxonomic Summary).

It is estimated that as little as 30-80 statistically independent Single Nucleotide Polymorphisms (SNP) can uniquely identify an individual human (15). The average sequence error rate (16) is greater than estimated human (intra-species) variation (17). Considering the poor coverage of unintended human content in the samples, even in the extreme lavage fluid examples, the total length of spot alignments identified as human are extremely unlikely to reveal validated, statistically independent SNPs capable of individual identification. The great majority of spots characterized by a human best hit though not the exclusive organism of the top five (“Conserved lineage spots” in Table 1) are highly significant alignments to related primates with approximately 20% sharing the same low *eValue* for all members (see Additional file 2, S1-S4). These likely represent conserved regions unfavored for SNP location (18).

## Conclusions

STAT has provided a successful framework for our SRA NGS submission pipeline. Sometimes actual sample content may be unknown, and submitted metadata are often incomplete and of poor quality (19, 20). Contamination, as highlighted above, may complicate or confuse further analysis. Recognizing these limitations stimulated our foremost goal to derive signals able to validate and accurately describe submitted data for the benefit of our users. Reflecting the National Institutes of Health (NIH) Science and Technology Research Infrastructure for Discovery, Experimentation, and Sustainability (STRIDES) Initiative (21), and ensuring that NIH-funded research data is findable, accessible, interoperable, and reusable (FAIR) (22), results from STAT are available through Amazon Web Services’ Athena and Google Cloud Platform’s BigQuery query services. Both can be searched to identify runs containing specific organismal content (23), allowing efficient selection of data for further analysis by the scientific community. Over approximately five years, we have processed more than 27.9 Peta base pairs from runs averaging 1.1 Giga base pairs in size with average total processing throughput of 3 minutes per run. While roughly 20% of runs analyzed to date are withheld by submitter request until ready for publication, nearly 10.8 million are publicly-queryable records, now richly annotated by STAT analysis.

Building a STAT database is flexible; it can be tailored to specific needs. For example, we are currently testing a STAT k-mer database designed to identify Antimicrobial Resistance (AMR) in NGS. The AMR_CDS FASTA file containing sequences curated by the NCBI Pathogen group (24) is used as input to generate 32 bp k-mers with a window size = 1; that is, the complete non-redundant k-mer set. For the purpose of removing human reads from clinical pathogen screening samples, we presented a tool combining STAT *aligns_to* with a human-specific database. As part of recent NIH-wide efforts to combat SARS-CoV-2, we released a detection tool containing *aligns_to* and a Virus “dbs” that allows users to map k-mers found in NGS data to taxa included under Coronaviridae (25). Our choice to maximize taxonomic coverage while minimizing k-mer count has proved a reasonable and effective balance. Employing the principle of MinHash in design, we contribute a framework others may find useful, and offer the collection of tools to use freely.

The success we and others have experienced is consistent with the notion of a random model of k-mer occurrence (26). Yet, as keenly shown by Breitwieser et al. (27) *unique* k-mer hits are the most informative. Through serendipity while preparing this manuscript, our colleague John Spouge enlightened us with his method of a non-parametric statistical approach to assess an NGS run using unique hits for confident measurement of taxonomic assignments^2^. We are just beginning to explore this implementation in STAT, and look forward to reporting results in the future.

## Methods

### General Design

STAT refers to a collection of tools for building k-mer databases, querying those databases, and reporting results of our SRA submission pipeline using the former. Details described below are based on our standard pipeline settings.

### k-mer Size

STAT uses 32 bp k-mers (i.e., k=32) for database generation, and as the unit for comparison. The majority of unaligned SRA data are reads between 60 and 150 bp in length, with mean error rate of 0.18% (16): such reads can be expected to yield many correct 32 bp k-mers for reliable identification. While reducing from 32 bp k-mers to 16 bp k-mers decreases the size of resulting databases, there is significant loss of specificity (10 ^9) per k-mer. By comparison, using 64 bp k-mers are extraordinarily more selective, but database size becomes impractical.

Finally, with each base encoded in 2 bits, 32 bp k-mers fit fully, and compactly in a 64-bit integer.

### k-mer Databases

Two primary types of k-mer databases are constructed (as described below): a dense database that selects one k-mer per 64 bp segment (“tree_filter”), of input sequence, while a sparse database (“tree_index”) selects one k-mer per 64 bp (Virus), 8000 bp (Eukaryota), and 2000 bp (Bacteria, and Archaea) segment respectfully, noting that segment size is roughly proportional to genome size.

### k-mer Generation

k-mers are selected using an iterative approach derived from MinHash (3). To compose STAT databases, for every fixed length segment (“window”) of incoming nucleotide sequence, a list of overlapping k-mers (effectively segment length plus right k-1 bp “wings”) is generated from both strands. The 32 bp k-mers are encoded using 2 bits per base into 64 bits (8 bytes), then a minimum k-mer representing this segment is selected based on the 64-bit encoded k-mer value read as 64-bit integer, effectively a 64-bit hash (see Figure 1).

### Taxonomic k-mer Database generation

Construction of k-mer databases is guided by the NCBI Taxonomy Database (5), specifically the four root Super Kingdoms: Archaea (722 species, 1330 total nodes), Bacteria (20,259 species, 29,835 total nodes), Eukaryota (455,421 species, 638,336 total nodes), and Viruses (4656 species, 7,583 total nodes) [current as of manuscript date (28)].

From each (Super Kingdom) root, lineage paths traverse nodes where terminal nodes are those containing only child leaves. Input sequences (see below) have an assigned NCBI Taxonomy Id (TaxId), and represent leaves on these trees. These lineage relationships are represented in a two-column file referred to as “parents”, wherein each node TaxId (first column) reports its parent node TaxId (second column).

All sequences (see Database Input Sequences) attached to a particular TaxId are input to k-mer database generation using segment (“window”) sizes as described. For each input set of sequences assigned a TaxId, the immediate output is a dictionary that contains the set of unique 32 bp k-mers derived as described (we designate this “db” file extension). Each dictionary is further transformed into a binary file that encodes every 32 bp k-mer as an 8-byte (64-bit) integer using 2 bits per base, followed by its TaxId represented in a 4-byte (32-bit) integer. Thus each k-mer record is stored as one 12-byte pair (k-mer, TaxId) in a database file designated with “dbs” file extension, sorted by k-mer for binary search optimization.

Next, using the taxonomic node relationships (found in the “parents” file), starting from the leaves we recursively merge each binary (“dbs”) file representing a unique set of k-mers derived from a single TaxId to sibling(s), then parent nodes. Each sibling leaf is merged such that k-mers specific to a leaf remain as diagnostic of that TaxId, while those found in neighboring (sibling species) leaves are moved up (“merged”) to the common parent node TaxId (see Figure 2). This process results in a single merged database file (“tree_filter.dbs”) representing all k-mers assigned a TaxId.

While it is difficult to generalize, we note that when the process of merging is complete, approximately 20% of the *Homo sapiens* 32 bp k-mers remain as unique to human; that is, 80% were not diagnostic for the species, and instead merged up the eukaryotic tree.

Database generation can be accomplished using any of the *build_index*\* tools (see github), and each takes parameters for window size, and k-mer size. The process of merging is accomplished using *merge_db*.

### Database Input Sequences

We use NCBI BLAST^®^ “refseq_genomes” database (29), supplemented with viral sequences extracted from the BLAST^®^ “nt/nr” database as input source for taxonomy identification in both sparse (“index”) and dense (“filter”) k-mer databases (30). Viral records are extracted from “nt/nr” by loading only sequences assigned a TaxId whose lineage root is the Super Kingdom “Viruses”.

### Querying the Taxonomic k-mer Database (STAT)

To query a k-mer database, input SRA accessions, or FASTA sequence is used to generate the unique set of query 32 bp k-mers read as 64-bit integer hashes for finding identical hashes (and assigned TaxId) from the designated k-mer database using the tool *aligns_to*. Proximal results are counts for each specific taxonomic k-mer hit (see Results, Figure 3). Passed an SRA accession, STAT built with NCBI NGS library support will retrieve query sequences, and *aligns_to* option *-unaligned_only* is available to limit analysis to the unaligned reads found in the SRA object.

### Database Filtering

We determined the need to delete low complexity k-mers composed of >50% homo-polymer or dinucleotide repeats (e.g. AAAAAA or ACACACACACA). This is accomplished using *filter_db*. We have also investigated “dusting” input sequences (31) and found it complementary to filtering, though it is not used at this time in our pipeline.

### Performance measurement

STAT performance metrics were gathered as described in Wood et al. (see “Execution of strain exclusion experiments”, and “Evaluation of accuracy in strain exclusion experiments” in Methods, 11). A “dense” k-mer database was created using the excluded taxa sequences for input (11). Briefly, we used Mason 2 (32) to generate 500,000 simulated Illumina 100 bp paired reads for each excluded strain TaxId, and collected cpu, and memory using ram-disk storage of the simulated reads employing 16 threads (16 Intel® Xeon® 2.8 GHz CPUs 64 GB RAM). Accuracy was measured using *aligns_to* against “tree_filter.dbss” (see Results) with a list of all TaxIds excluding the 50 strains tested (130,769 TaxIds total).

### SARS-CoV-2 Contamination Identification and Verification

Submissions suggesting contamination with SARS-CoV-2 identified through normal processing were subject to two further verification methods. All identified accessions were rerun using the current SARS-CoV-2 Detection tool (25, *DockerHub Tag 1*.*1*.*2021-01-25*, see Additional file 1). Low level contamination (1 spot, 1 or 0 resolved hits) observed in 31 records was further examined using STAT against a SARS-CoV-2 - specific database (“dbs”) composed of 32-bp kmers identified by Wahba et al. (33). Using these 18582 SARS-CoV-2 - specific k-mers as queries never found a matching k-mer when run against our full tree_filter.dbs (data not shown).

### Human Contamination Identification and Removal

The special-purpose k-mer database uses NCBI BLAST^®^ “refseq_genomes” limited to Human (TaxId 9606) for input using a “window” segment of 32 bp and filtered as described previously. Any k-mers found also in the merged Kingdom databases of Bacteria, and Viruses were removed. The current database contains 80,143,408 k-mers and is 612 mb in size. The *sra-human-scrubber* is intended as the last step before submission and takes as input a “fastq file”, and outputs a “fastq.clean file” in which all reads identified as potentially of human origin are removed (34).

Examples discussed in Results and shown in Table 1 were run against the *sra-human-scrubber* docker container ( 34, *DockerHub Tag 1*.*0*.*2021-03-11*). For each, the resulting “{file}.fastq.clean” was transformed into a fasta file, and then subject to NCBI *blastn* 2.10.0+ using (megablast) parameters [*-max_target_seqs* 5, -*evalue* 0.00001, *-strand* plus] against the “refseq_genomes” BLAST^®^ database (35). The top five hits (by *eValue*) for each spot containing a human best hit (with all hits *eValue* < 1e -10) can be found in Additional file 2.

## Additional Files

- *Additional file 1*.*xlsx*, Microsoft Excel: The first sheet (S1) contains results from accessions using SARS-CoV-2 detection tool as described in Methods; the second sheet (S2) contains those accessions from S1 subject to verification using STAT as described in Methods.

- *S1 SARS-CoV-2 Contamination*
- *S2 SARS-CoV-2 Verification*
- *Additional file 2*.*xlsx*. Microsoft Excel: The first four sheets (S1-S4) contain the top five NCBI BLAST^®^ hits for each accession spot in which at least one of those hits was human. The last sheet contains summary STAT taxonomic data for each of the four accessions before and after human contamination removal tool treatment as described in Methods.

- *S1 SRR11092056 BLAST*^®^*Results*
- *S2 SRR11092057 BLAST*^®^ *Results*
- *S3 SRR13402847 BLAST*^®^ *Results*
- *S4 SRR13444106 BLAST*^®^ *Results*
- *S5 STAT Taxonomic Slices*

## Declarations

### Ethics approval and consent to participate

- Not applicable

### Consent for publication

- Not applicable

### Availability of data and materials

- https://github.com/ncbi/ngs-tools/tree/tax/tools/tax/src
- https://hub.docker.com/r/ncbi/sra-human-scrubber
- https://hub.docker.com/r/ncbi/SARS-CoV-2-detection-tool

### Authors’ contributions

- KSK prepared the manuscript including figures, executed performance and accuracy analyses, as well as generated all supplementary data.
- OS authored most of the C++ / python STAT tools.
- JRB, CO contributed to discussions of design, and implementation.
- RL tested early versions of the tools.

## Acknowledgements

- Vadim Zalunin, Alex Efremov, and Andrey Kochergin for building, maintaining, and improving the STAT pipeline.
- Ryan Connor for always stimulating conversation.
- Christiam Camacho for generous NCBI BLAST^®^ support.
- David Lipman for ideas, and (always) vigorous discussion.
- Benjamin Langmead for supplying strain exclusion fasta files.
- Susan Roberts and Lydia Fleischmann for indispensable editing assistance.

## Footnotes

1 We use the word “spot” to reference either the un-split paired biological read, or the single unpaired biological read.

2 John Spouge, Statistical Computational Biology Group, National Library of Medicine, National Institutes of Health, Personal communication.

## References

1. Shumway M, Cochrane G, Sugawara H. Archiving next generation sequencing data. Nucleic Acids Res. 2010 Jan;38 Database issue:D870–1. Available from: DOI: 10.1093/nar/gkp1078

2. Kodama Y, Shumway M, Leinonen R; International Nucleotide Sequence Database Collaboration. The Sequence Read Archive: explosive growth of sequencing data. Nucleic Acids Res. 2012 Jan;40 Database issue:D54–6. Available from: DOI: 10.1093/nar/gkr854

3. Broder AZ. Identifying and filtering near-duplicate documents. In: COM ‘00 Proceedings of the 11th Annual Symposium on Combinatorial Pattern Matching. London: Springer; 2000; 1848;p.1–10. Available from: https://doi.org/10.1007/3-540-45123-4_1

4. Ondov, B.D., Treangen, T.J., Melsted, P. et al. Mash: fast genome and metagenome distance estimation using MinHash. Genome Biol. 2016;17:132. Available from: https://doi.org/10.1186/s13059-016-0997-x

5. NCBI Taxonomy Browser [internet].Taxonomy [Internet]. Available from: https://www.ncbi.nlm.nih.gov/taxonomy/

6. Brister JR, Ako-Adjei D, Bao Y, Blinkova O. NCBI viral genomes resource. Nucleic Acids Res. 2015 Jan;43(Database issue):D571–7. Available from: DOI: 10.1093/nar/gku1207

7. Simmonds P, Adams MJ, Benkő M, Breitbart M, Brister JR, Carstens EB, et al. Consensus statement: Virus taxonomy in the age of metagenomics. Nat Rev Microbiol. 2017 Mar;15(3):161–168. Available from: DOI: 10.1038/nrmicro.2016.177

8. A sea change for virology. Nat Rev Microbiol. 2017 Feb 13;15(3):129. Available from: DOI: 10.1038/nrmicro.2017.13

9. Breitwieser FP, Lu J, Salzberg SL. A review of methods and databases for metagenomic classification and assembly. Brief Bioinform. 2019;20:1125– 1136. Available from: DOI: 10.1093/bib/bbx120

10. Wood DE, Salzberg SL. Kraken: ultrafast metagenomic sequence classification using exact alignments. Genome Biol. 2014;15:R46. Available from: DOI: 10.1186/gb-2014-15-3-r46

11. Wood DE, Lu J, Langmead B. Improved metagenomic analysis with Kraken 2. Genome Biol. 2019 Nov 28;20(1):257. Available from: DOI: 10.1186/s13059-019-1891-0

12. Al-Qahtani AA. Severe Acute Respiratory Syndrome Coronavirus 2 (SARS-CoV-2): Emergence, history, basic and clinical aspects. Saudi J Biol Sci. 2020 Oct 27(10):2531–2538. Available from: DOI: 10.1016/j.sjbs.2020.04.033

13. Lillie PJ, Samson A, Li A, Adams K, Capstick R, Barlow GD et al. Novel coronavirus disease (Covid-19): The first two patients in the UK with person to person transmission. J Infect. 2020 May;80(5):578–606. Available from: DOI: 10.1016/j.jinf.2020.02.020

14. Shabani M, Marelli L. Re-identifiability of genomic data and the GDPR: Assessing the re-identifiability of genomic data in light of the EU General Data Protection Regulation. EMBO Rep. 2019 Jun;20(6):e4831. Available from: DOI: 10.15252/embr.201948316

15. Lin Z, Owen AB, Altman RB. Genetics. Genomic research and human subject privacy. Science. 2004 Jul 9;305(5681):183. Available from: DOI: 10.1126/science.1095019

16. Pfeiffer F, Gröber C, Blank M, Händler K, Beyer M, Schultze JL, et al. Systematic evaluation of error rates and causes in short samples in next-generation sequencing. Sci Rep. 2018 Jul 19;8(1):10950. Available from: 10.1038/s41598-018-29325-6

17. Chakravarti A. Perspectives on Human Variation through the Lens of Diversity and Race. Cold Spring Harb Perspect Biol. 2015 Sep 1;7(9):a023358. Available from: DOI: 10.1101/cshperspect.a023358

18. Castle JC. SNPs occur in regions with less genomic sequence conservation. PLoS One. 2011;6(6):e20660. DOI: 10.1371/journal.pone.0020660

19. Bernstein MN, Doan A, Dewey CN. MetaSRA: normalized human sample-specific metadata for the Sequence Read Archive. Bioinformatics. 2017 Sep 15;33(18):2914–2923. Available from: DOI: 10.1093/bioinformatics/btx334

20. Bernstein MN, Gladstein A, Latt KZ, Clough E, Busby B, Dillman A. Jupyter notebook-based tools for building structured datasets from the Sequence Read Archive. F1000Res. 2020 May 19;9:376. Available from: https://doi.org/10.12688/f1000research.23180.2.

21. NIH Office of Data Science Strategy [internet]. STRIDES. Available from: https://datascience.nih.gov/strides

22. Wilkinson MD, Dumontier M, Aalbersberg IJ, Appleton G, Axton M, Baak A, et al. The FAIR Guiding Principles for scientific data management and stewardship. Sci Data. 2016 Mar 15;3:160018. Available from: DOI: 10.1038/sdata.2016.18

23. NCBI Sequence Read Archive (SRA) [internet]. SRA in the cloud. Available from: https://www.ncbi.nlm.nih.gov/sra/docs/sra-cloud-based-examples/

24. NCBI National database of antibiotic resistant organisms (NDARO). AMR cds fasta. Available from: https://ftp.ncbi.nlm.nih.gov/pathogen/Antimicrobial_resistance/AMRFinderPlus/data/latest/AMR_CDS

25. NCBI Sequence Read Archive (SRA) [internet]. SRA detection tool. Available from: https://www.ncbi.nlm.nih.gov/sra/docs/sra-detection-tool

26. Fofanov Y, Luo Y, Katili C, Wang J, Belosludtsev Y, Powdrill T, et al. How independent are the appearances of n-mers in different genomes? Bioinformatics. 2004 Oct 12;20(15):2421–8. Available from: DOI: 10.1093/bioinformatics/bth266

27. Breitwieser FP, Baker DN, Salzberg SL. KrakenUniq: confident and fast metagenomics classification using unique k-mer counts. Genome Biol. 2018;19:198. Available from: DOI: 10.1186/s13059-018-1568-0

28. NCBI Taxonomy Browser.Taxonomy Statistics [Internet]. Taxonomy Nodes (all dates). Available from: https://www.ncbi.nlm.nih.gov/Taxonomy/taxonomyhome.html/index.cgi?chapter=statistics&uncultured=hide&unspecified=hide

29. Pruitt KD, Tatusova T, Brown GR, Maglott DR. NCBI Reference Sequences (RefSeq): current status, new features and genome annotation policy. Nucleic Acids Res. 2012 Jan;40(Database issue):D130–5. Available from: DOI: 10.1093/nar/gkr1079

30. NCBI FTP [Internet]. The BLAST® Databases. Available from: https://ftp.ncbi.nlm.nih.gov/blast/documents/blastdb.html

31. Morgulis A, Gertz EM, Schäffer AA, Agarwala R. A fast and symmetric DUST implementation to mask low-complexity DNA sequences. J Comput Biol. 2006 Jun;13(5):1028–40. Available from: DOI: 10.1089/cmb.2006.13.1028

32. Holtgrewe, M. Mason - a read simulator for second generation sequencing data. Technical Report 2010. Available from: DOI: 10.17169/refubium-22374

33. Wahba L, Jain N, Fire AZ, Shoura MJ, Artiles KL, McCoy MJ, Jeong DE. An Extensive Meta-Metagenomic Search Identifies SARS-CoV-2-Homologous Sequences in Pangolin Lung Viromes. mSphere. 2020 May 6;5(3):e00160–20. Available from: DOI: 10.1128/mSphere.00160-20

34. Docker Hub [internet]. NCBI sra-human-scrubber Docker image. Available from: https://hub.docker.com/r/ncbi/sra-human-scrubber

35. Morgulis A, Coulouris G, Raytselis Y, Madden TL, Agarwala R, Schäffer AA. Database indexing for production MegaBLAST searches. Bioinformatics. 2008 Aug 15;24(16):1757–64. Available from: DOI: 10.1093/bioinformatics/btn322

